# Chronic Treatment of a Mouse Model of Cerebral Amyloid Angiopathy and Brain AT_1_ Receptor Expression

**DOI:** 10.1101/2025.05.16.654535

**Authors:** Natalia Motzko Noto, Lisa S. Robison, Robert C. Speth

## Abstract

**Introduction:** The renin-angiotensin-aldosterone system (RAAS) has been shown to be dysregulated in dementia, with elevated levels of angiotensin-converting enzyme (ACE), angiotensin (Ang) II, and Ang II type 1 receptors (AT_1_Rs). Cerebral amyloid angiopathy (CAA), a common cerebrovascular disease, currently has no treatment or cure available. We aimed to determine if a mouse model with CAA (Tg-SwDI) also exhibits elevated levels of AT_1_Rs and whether RAAS-targeting drugs (telmisartan and lisinopril) mitigate these effects.

**Materials and Methods:** Tg-SwDI mice were treated with sub-depressor doses of either telmisartan or lisinopril from 3-8 months of age, with blood pressure being monitored 2 and 4 months after the start of treatment. Postmortem, receptor autoradiography was performed to determine levels of AT_1_R in 13 brain regions in untreated and treated Tg-SwDI mice compared to wild-type controls (C57Bl/6J).

**Results:** No statistically significant differences among groups were observed in any of the 13 regions analyzed. However, trends with medium to large effect sizes were observed.

**Conclusions:** CAA did not significantly dysregulate AT_1_R levels in the brains of Tg-SwDI mice compared to wild-type mice. Drug treatment caused no significant brain AT_1_R alterations. Further studies are required to determine if the trends observed are pathophysiological and pharmacologically significant.

## 1. Introduction

### 1.1. The renin-angiotensin-aldosterone system (RAAS) and RAAS-targeting drugs

The RAAS is widely known to play a role in blood pressure regulation, fluid homeostasis, electrolyte balance, and systemic vascular resistance^1–3^. It is divided into a classical arm and a counterregulatory arm. Activation of the classical angiotensin-converting enzyme (ACE)/angiotensin (Ang) II/Ang II receptor type 1 (AT_1_R) arm causes vasoconstriction, inflammation, hypertrophy, cellular proliferation, and salt and water reabsorption; while activation of the counterregulatory ACE-2/Ang-(1-7)/Mas receptor (MasR) arm counteracts Ang II signaling through AT_1_Rs^4^. Of note, Ang II can activate the AT_2_R, which also counteracts the AT_1_R^5^ (**Figure 1A**).

**Figure 1.**
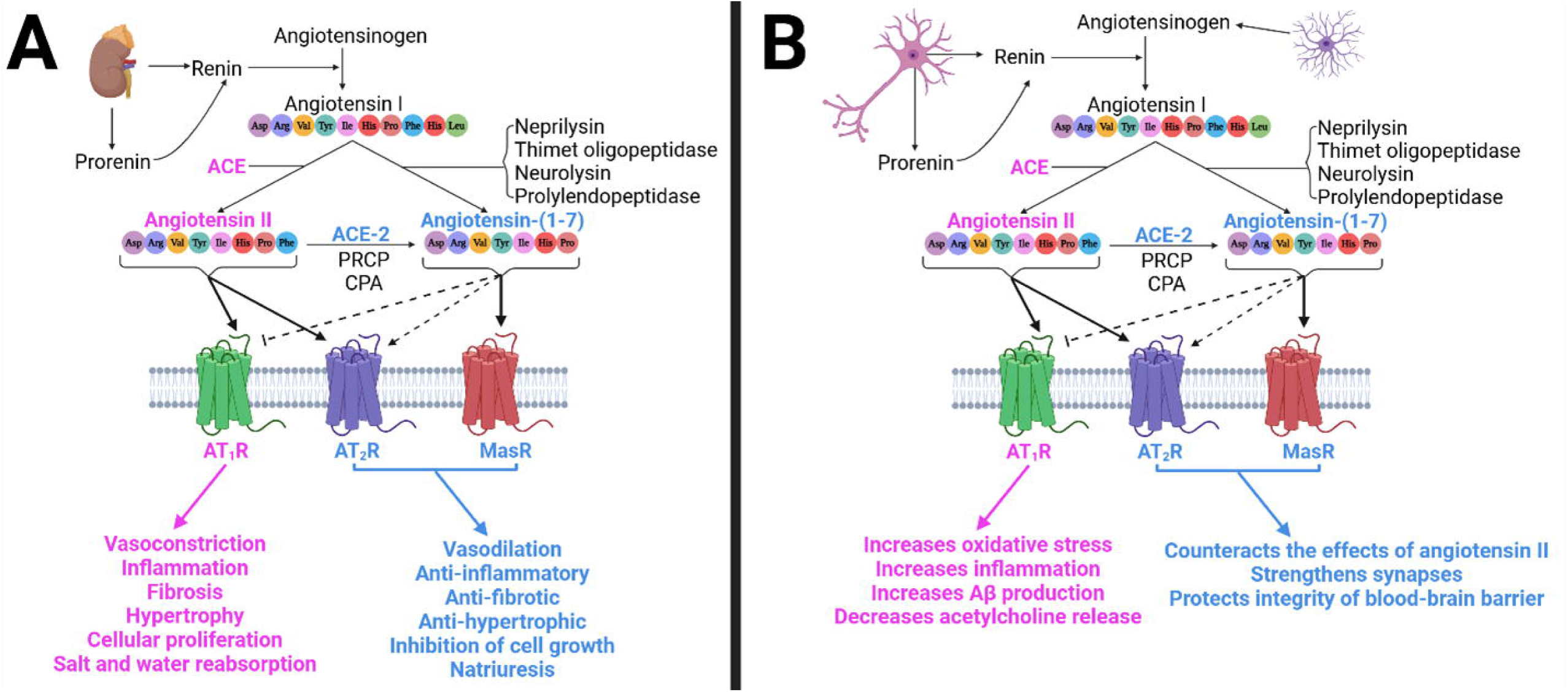
Simplified representation of the RAAS activity in the vasculature and in the brain. **(A)** In the vasculature, the kidneys release the enzyme renin, along with its precursor prorenin. Renin is responsible for cleaving angiotensinogen, giving rise to angiotensin (Ang) I, which is then cleaved by the angiotensin-converting enzyme (ACE), generating Ang II. In turn, Ang II can activate both Ang II type 1 receptors (AT_1_Rs) and Ang II type 2 receptors (AT_2_Rs). Additionally, Ang II can be further cleaved by ACE-2, prolylcarboxypeptidase (PRCP), or carboxypeptidase A (CPA) to generate Ang-(1-7), which can activate MasRs and AT_2_Rs (to a lesser extent), and inhibit AT_1_Rs, counteracting the effects of Ang II. Alternatively, Ang-(1-7) can also be formed through cleavage of Ang I by different enzymes. **(B)** In the brain, astroglia produce angiotensinogen^128^, which can be cleaved by renin, prorenin, or a renin-like enzyme present in neurons^129^, to give rise to Ang I^130, 131^. In turn, ACEs, which are expressed on the cell membranes of the brain, cleave Ang I, giving rise to Ang II^132^. Like in the vasculature, Ang-(1-7) activity in the brain counteracts the effects of Ang II. *Created in BioRender. Speth, R. (2025)* https://BioRender.com/rzfj0v4

Physiologically, renin is produced in response to low blood pressure, activating Ang II to increase vascular resistance and blood pressure; however, hyperactivation of the classical RAAS arm results in chronic hypertension and other pathological conditions. Drugs targeting the RAAS, such as renin inhibitors, ACE inhibitors, and angiotensin receptor blockers (ARBs) have been developed to reverse hypertension^6, 7^. These drugs prevent overactivation of the RAAS by inhibiting renin and ACE activity to decrease Ang II formation; or by blocking the AT_1_R, leaving Ang II to either bind to the AT_2_R or to be converted to Ang-(1-7).

### 1.2. The RAAS system in the brain

The RAAS also exists in brain, impacting cardiovascular control and neurological processes, such as cognition and anxiety^8, 9^. Ang II activates both AT_1_Rs and AT_2_Rs in the brain. Binding to the AT_1_Rs, found on neurons, astrocytes, oligodendrocytes, and microglia present in the cortex, hippocampus, and basal ganglia, causes inflammation, oxidative stress, and cell death, contributing to cognitive impairment^10^. Conversely, Ang II binding to the AT_2_Rs has antioxidant and anti-inflammatory properties, enhancing cell survival and supporting cognitive performance^10^ (**Figure 1B**). Additionally, Ang-(1-7) reportedly interacts with the Mas receptors (MasRs) in brain areas such as the hippocampus and cerebral cortex^11, 12^, reportedly strengthening synapses involved in memory^8^.

Another key mechanism by which the RAAS can influence brain health is by impacting cerebrovascular integrity and cerebral blood flow. Even under resting conditions, the brain requires 15-20% of cardiac output to sustain its metabolic activity^13, 14^ and properly clear metabolic byproducts^15^. Hypertension challenges the integrity of the cerebral vasculature, causing endothelial dysfunction, vascular remodeling, and hypoperfusion^16^, leading to microvascular rarefaction and dysfunction and neurovascular uncoupling, impairing cerebral blood flow^15, 17^.

### 1.3. The RAAS as a target for dementia treatment

Dysregulation of the RAAS, both systemically and centrally, is associated with neurodegenerative diseases including dementia, characterized by impairment in thinking, reasoning, and memory. Two of the most common forms of dementia are Alzheimer’s disease (AD), characterized by amyloid-beta (Aβ) plaques and tau tangles, and vascular cognitive impairment and dementia (VCID), which includes cardiovascular and cerebrovascular disease-related cognitive decline. Hypertension, especially in mid-life, is a major risk factor for dementia^15, 17, 18^. Additionally, the brain RAAS is dysregulated in dementias such as AD, with previous studies in humans and mouse models of dementia showing elevated levels of ACE, Ang II, and AT_1_Rs^19–21^.

Over the years, efforts have been made to repurpose RAAS-targeting drugs to prevent or treat neurodegenerative diseases^22–24^. This comes after evidence that the use of antihypertensive drugs, particularly those that can cross the blood-brain barrier (BBB), are associated with a reduced risk of dementia, independent of blood pressure alteration^25–27^. Studies showed that centrally-acting ACE inhibitors (mainly captopril) can retard the development of neurodegeneration and even reduce cognitive decline^25, 28^, while non-centrally-acting ACE inhibitors reportedly increase the risk of dementia by 73% in older hypertensive adults^25^.

#### 1.3.1. Alzheimer’s disease (AD)

Antihypertensive drug treatment reduces the risk of AD. A case-control study conducted in Italy on patients who were 65 or older, between 2009 and 2012, showed a significant inverse association between exposure to antihypertensive drug treatment and AD^29^. Notably, antihypertensive drugs that target the RAAS specifically, appear to have neuroprotective benefits above and beyond their ability to control blood pressure. A secondary longitudinal data analysis of normally cognitive adults who were 75 or older, showed that usage of ACE inhibitors and ARBs was associated with reduced risk of AD^30^. Additionally, studies performed in an AD mouse model (ddY mouse that is intracerebroventricularly injected with Aβ) showed that treatment with the ARB telmisartan reduced Aβ deposition in the brain, consequently reducing the rate of cognitive decline^31, 32^.

#### 1.3.2. Vascular cognitive impairment and dementia (VCID)

A systematic review of 38 relevant publications ranging from longitudinal studies, to randomized controlled trials, to meta-analyses, concluded that antihypertensive medications reduced the risk of both vascular dementia and cognitive decline^33^. Additionally, a regression analysis showed that ARBs in particular can reduce the risk of dementia with micro-vascular origin^34^.

#### 1.3.3. Stroke

RAAS-targeting drugs are also effective against other cerebrovascular conditions, such as stroke. In the early 2000s, the HOPE study^35^ found that ramipril, a centrally-acting ACE inhibitor, reduced the risk of stroke by 32% and the risk of fatal stroke by 61%, compared to placebo, despite modest reductions in blood pressure^36^. Similarly, a double-masked, randomized, parallel-group trial of the ARB losartan in hypertensive patients 55-80 years old reduced the rate of stroke (fatal and non-fatal) by 25% more than an equi-antihypertensive dose of the beta blocker, atenolol ^37^.

### 1.4. Cerebral amyloid angiopathy (CAA)

CAA is a common cerebrovascular disease^38^ caused by Aβ protein accumulation in the tunica media layer of the walls of cerebral arteries, arterioles, and capillaries of the leptomeninges and cortex, usually surrounding smooth muscle cells^38–42^. It causes perivascular inflammation due to elevated levels of reactive astrocytes and activated microglia around the vessels as the disease progresses^43^. CAA causes VCID, associated with specific cognitive impairments (e.g., perceptual speed and episodic memory) ^43, 44^.It is also associated with increased stroke risk^45^. As it progresses, pathological changes in the blood vessels occur, including destruction of smooth muscle cells in the tunica media, replacement of tunica media with Aβ, microaneurysm formation, fibrinoid necrosis, and cerebral hemorrhage^39, 46^. Moreover, CAA pathology is found in over 90% of AD brains^47^; both conditions involve Aβ accumulation, although the location of where it accumulates differs. In the presence of AD, CAA exacerbates AD pathology and symptoms^40, 47^. Additionally, the presence of CAA in the AD brain may preclude these patients from receiving treatment with anti-amyloid monoclonal antibodies due to an increased risk of amyloid-related imaging abnormality (ARIA), including brain swelling (ARIA-E) and/or bleeding (ARIA-H) ^48–50^.

Currently, there are no pharmaceutical agents available to cure or treat CAA, and development of new treatments is proving to be challenging^47, 51^, due to the high rate of meningoencephalitis, increased risk of ARIA-E and ARIA-H, or simply lack of efficacy^51^.

### 1.5. Premise of study

As mentioned previously, there is a dysregulation of the RAAS in the AD brain^19–21^, suggesting there may be potential for repurposing RAAS-targeting drugs to treat neurodegenerative diseases, particularly AD^52, 53^. To the best of our knowledge, no studies have determined the role that the RAAS might have in CAA, or the role of RAAS-targeting drugs to mitigate CAA.

In this study, we hypothesized that RAAS-targeting drugs would alter the expression of AT_1_R in the brains of Tg-SwDI mice, that brain AT_1_R expression would differ between Tg-SwDI and wild-type mice, and that there would be sex differences in brain AT_1_R expression. We investigated whether AT_1_R binding was altered in Tg-SwDI mice, a mouse model exhibiting CAA, across several brain regions. Additionally, we treated Tg-SwDI mice with RAAS-targeting drugs from two different classes, telmisartan (ARB) and lisinopril (ACE inhibitor), during early-stage disease to assess their ability to mitigate changes in brain AT_1_R expression that might be associated with CAA.

## 2. Materials and Methods

### 2.1. Animals

Male and female Tg-SwDI mice (JAX MMRRC Stock # 034843) served as a transgenic animal model of CAA; this strain exhibits behavioral deficits, fibrillar vascular Aβ pathology, and perivascular inflammation^54^. In this model, Aβ accumulates primarily around the cerebral vasculature, beginning at approximately 3-months of age, with severity increasing over the lifespan^55–57^. At 12-months of age, the hippocampus is most affected by CAA, followed by the thalamus^54^. However, CAA is almost absent in the cortical vasculature^54^. Age-and sex-matched C57BL/6J mice, the background strain of Tg-SwDI mice, served as wild-type (WT) controls (*N*=105, divided across 3 consecutive cohorts).

Males and females were kept in separate cages, with 3-to-5 littermates, on a reverse 12-hour light cycle (lights on at 8/9 PM, lights off at 8/9AM, varying with daylight-savings time). Temperature and humidity were 68-72°F and 40-60%, respectively. Food and water were available ad libitum.

The Nova Southeastern University Institutional Animal Care and Use Committee approved the experimental procedures used in this study (approval no. 2022.04.LR2). All animal housing and experiments were conducted in strict accordance with the institutional Guidelines for Care and Use of Laboratory Animals at Nova Southeastern University.

A timeline of the study is shown in **Figure 2**.

**Figure 2.**
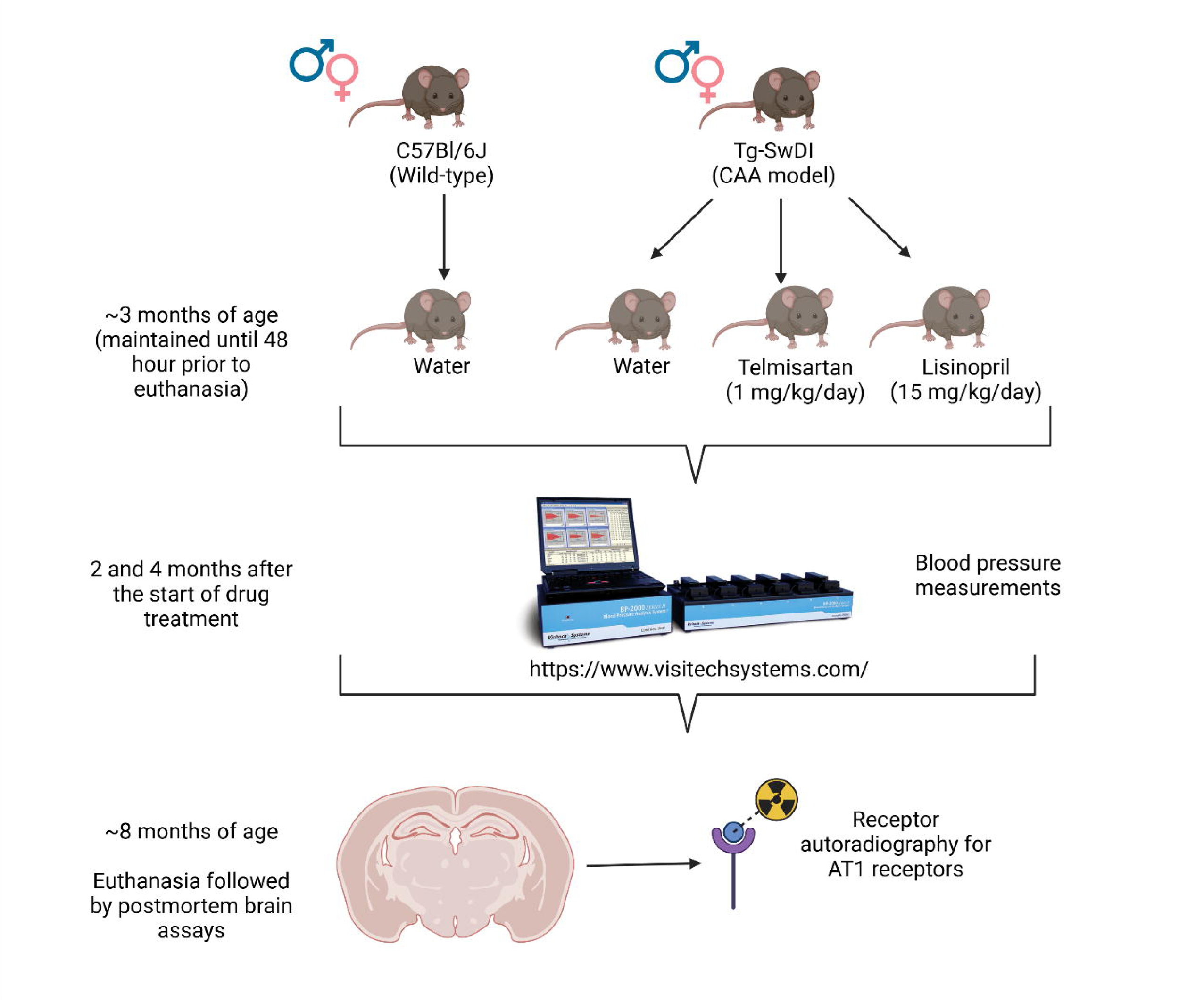
Generic timeline of the study for each cohort. Mice started on the drug treatment at approximately 3 months of age, undergoing blood pressure measurements 2 and 4 months into treatment. Euthanasia occurred at approximately 8 months of age, when significant CAA pathology is observed in Tg-SwDI mice, followed by cryostat sectioning of brains and RAR. *Created in BioRender. Speth, R. (2025)* https://BioRender.com/t7lrdza

### 2.2. Drug treatments

Tg-SwDI mice underwent a 5-month-drug treatment, starting at approximately 3-months of age, prior to the onset of significant Aβ pathology^55, 56^. They were euthanized for tissue collection at approximately 8-months of age, when significant CAA pathology and cognitive impairments appear in this animal model^58^.

Mice assigned to drug groups received sub-depressor doses of either telmisartan (1 mg/kg/day^59^) or lisinopril (15 mg/kg/day^60, 61^) dissolved in reverse osmosis water. Telmisartan solutions were made by diluting the powdered drug in 1-2 mL of 0.1 M sodium hydroxide, then mixing it with reverse osmosis water to achieve calculated concentration. pH was adjusted to 7.0-7.4 with 0.1 M hydrochloric acid. Solutions were covered in aluminum foil at all times due to the light-sensitive properties of telmisartan^62^. Lisinopril solutions were made by diluting powdered drug in reverse osmosis water to reach calculated concentration. Additional groups of control Tg-SwDI and C57Bl/6J mice received plain drinking water only. **Table 1** shows groups and sample sizes.

**Table 1.** Mouse distribution per cohort.

| **<u>Strain</u>** | **<u>Sex</u>** | **<u>Group</u>** | **<u>N</u>**<br>**<u>(2-month BP)</u>** | **<u>N</u>**<br>**<u>(4-month BP)</u>** | **<u>N</u>**<br>**<u>(RAR)</u>** |
| --- | --- | --- | --- | --- | --- |
| WT (C57BL/6J) | Male | Water | 14 | 14 | 5 |
| WT (C57BL/6J) | Female | Water | 14 | 14 | 5 |
| Tg-SwDI | Male | Water | 15 | 15 | 5 |
| Tg-SwDI | Female | Water | 12 | 12 | 4 |
| Tg-SwDI | Male | Telmisartan | 12 | 12 | 4 |
| Tg-SwDI | Female | Telmisartan | 13 | 13 | 5 |
| Tg-SwDI | Male | Lisinopril | 14 | 12 | 5 |
| Tg-SwDI | Female | Lisinopril | 14 | 14 | 5 |

\*Table shows how many animals participated in BP measurements and RAR but does not reflect outliers removed from analysis by Grubbs' test.

### 2.3. Blood pressure (BP) measurements

To ensure that the drug doses were in fact sub-depressor, BP was measured via the tail-cuff method^63, 64^ at 2 (*N*=108; 1 outlier removed) and 4 (*N*=106; no outliers) months after start of drug treatment using a BP-200 Blood Pressure Analysis System (Visitech Systems). BP measurements were performed for 4 consecutive days, with day 1 being an acclimation day to obtain reliable measurements. Each day, 7 preliminary measurements were taken to acclimate the mice to the procedure, followed by 10 replicate measurements that were averaged out for each mouse’s daily average BP.

### 2.4. Tissue collection

Drug treatment was suspended 48 hours prior to euthanasia to prevent any residual telmisartan from blocking AT_1_R binding. During this period, mice received plain water. Mice were deeply anesthetized with isoflurane anesthesia (5% induction; ∼3-4% maintenance) and perfused intracardially with chilled 0.9% saline prior to decapitation and collection of the brain. Brains were flash frozen in isopentane (*N*=105) and stored at-80°C until processing.

### 2.5. Brain sectioning

Brains were cut coronally at a thickness of 20-microns using a cryostat. Sections were thaw-mounted onto microscope slides (Fisherbrand^TM^ Tissue Path Superfrost^TM^ Plus Gold Slides), starting at approximately 1.10 mm rostral to Bregma and ending at approximately-2.91 mm caudal to Bregma^65^. Sections were mounted on 3 sets of slides: one pre-hippocampal set with 6 slides of ∼12 sections, and two post-hippocampal sets of 8 slides each with ∼6 sections. Once slide-mounted, sections were left to air-dry for approximately 30 minutes before being stored at-20°C until used for the receptor autoradiography protocol.

### 2.6. Receptor autoradiography (RAR)

RAR was performed on a subset of brains to determine the concentration of AT_1_R within different brain regions, following a previously described procedure^66^. The first 2 slides of each set were used. The first (-1) slides for each mouse were used to determine non-AT_1_R binding of ^125^I-sarcosine^1^,isoleucine^8^ Ang II (^125^I-SI Ang II) prepared as described previously^67^ [nonspecific binding (NSP)] in the presence of 10 µM losartan (a specific AT_1_R antagonist), while the second (-2) slides for each mouse were used to determine total ^125^I-SI Ang II binding in the absence of losartan. The assay was carried out in glass Coplin jars divided into 3 steps: pre-incubation, incubation, and rinses (**Figure 3**). Once slides were fully dry, they were taped to cardboards, along with an ^125^Iodine calibration standard slide^66^. Cardboards were placed inside X-ray cassettes with single-sided emulsion film (Carestream, MR-1). Cassettes were kept at-20°C for 4-7 days, after which the films were developed using D-19 type developer (Photographers’ Formulary, Condon, MT, USA, Catalog # 01-0035). Films were left to air-dry before being scanned at 2,400 dpi resolution in a Cannon scanner (Canoscan LIDE 700F).

**Figure 3.**
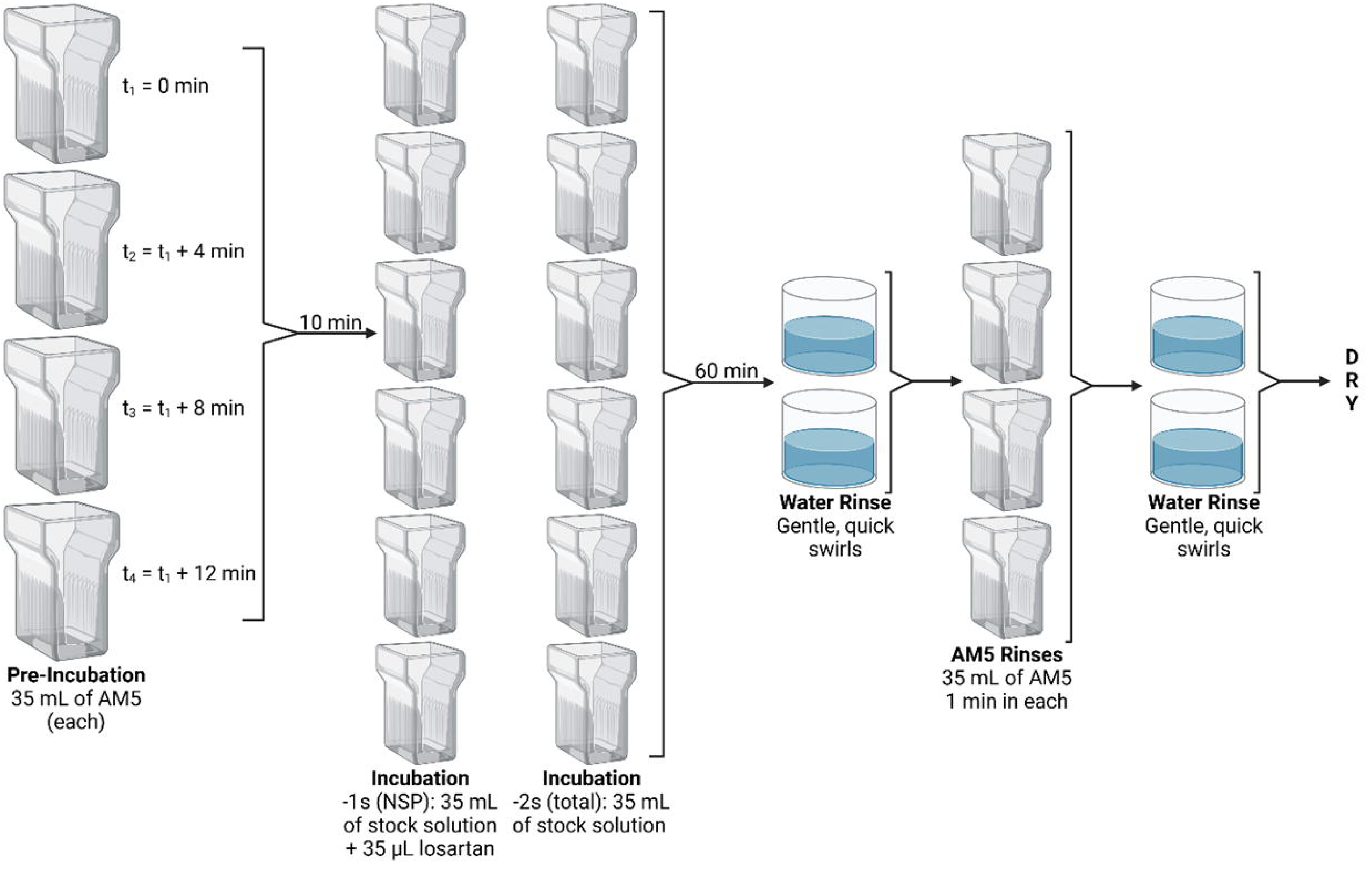
RAR experimental design. Four Coplin jars that can hold 10 slides each were filled with AM5 and used to carry out the pre-incubation step. After 10 min in solution, slides were immediately transferred to their respective incubation jars, where they incubated for 60 min at room temperature. Each incubation jar contained 35 mL of a stock solution prepared by mixing assay medium 5 (AM5) solution (150 mM sodium chloride, 5 mM disodium EDTA, 0.1 mM Bacitracin, and 500 mM dibasic sodium phosphate; pH 7.1 – 7.2) and ^125^I-SI Ang II ligand to achieve a concentration of 255 pM. Additionally, 35 μL of 10 μM losartan were added to half of the Coplin jars. Slides were blotted using a paper towel before the first two water rinses and after the last two water rinses. Each water rinse swirl did not last for more than 1 or 2 seconds. *Created in BioRender. Speth, R. (2025)* https://BioRender.com/omwatj2

Scanned films were used to sample brain regions of interest (ROIs) using ImageJ software to determine the concentration of total and non-AT_1_R (NSP) binding in fmol/mg initial wet weight. Specific AT_1_R binding was calculated by subtracting the concentrations obtained for non-AT_1_R (NSP) binding from that of total binding.

### 2.7. Statistical analysis

Data were analyzed using 2-way ANOVAs (factors: strain/treatment and sex), followed by Tukey’s multiple comparisons post-hoc tests. Outliers were identified by Grubbs’ test and removed prior to analyses. Threshold for statistical significance was set at p<0.05, while trends were described when p<0.10. Partial eta squared (η_p_ ^2^) was calculated to determine the effect size of main effects (treatment and sex differences). Cohen’s d (d) was calculated to determine the effect size of multiple pairwise comparisons tests. Results are presented as mean±SEM. GraphPad Prism Version 10.3.1 (GraphPad Prism, San Diego, CA) was used for all data analyses and graphing.

## 3. Results

### 3.1. Drugs at target doses did not alter systolic BP of treated versus untreated Tg-SwDI mice Drug intake was calculated by cage in mg/kg BW/day on a weekly basis and averaged

across the study. A one-sample t-test was used to determine whether target doses were reached, assessed by a lack of statistical significance between average drug intake and the target doses (1 mg/kg BW/day and 15 mg/kg BW/day for telmisartan and lisinopril, respectively). Both drugs did not alter BP in either males or females either after 2 or 4 months of treatment (**Figure 4A-C**). Tg-SwDI mice, regardless of treatment group, exhibited significantly lower systolic BP than WTs at 2 and 4 months treatment. When separated by sex, only telmisartan-treated Tg-SwDI male mice had significantly lower BP than WTs at 2 months, while lisinopril-treated males and females had significantly lower BP than WTs at 4 months. In females, all Tg-SwDI groups exhibited significantly lower systolic BP compared to WTs at 2 months. Overall, males displayed higher systolic BP than females; when divided into treatment groups, this sex difference was maintained across all Tg-SwDI groups at 2 months, and all 4 groups at 4 months.

**Figure 4.**
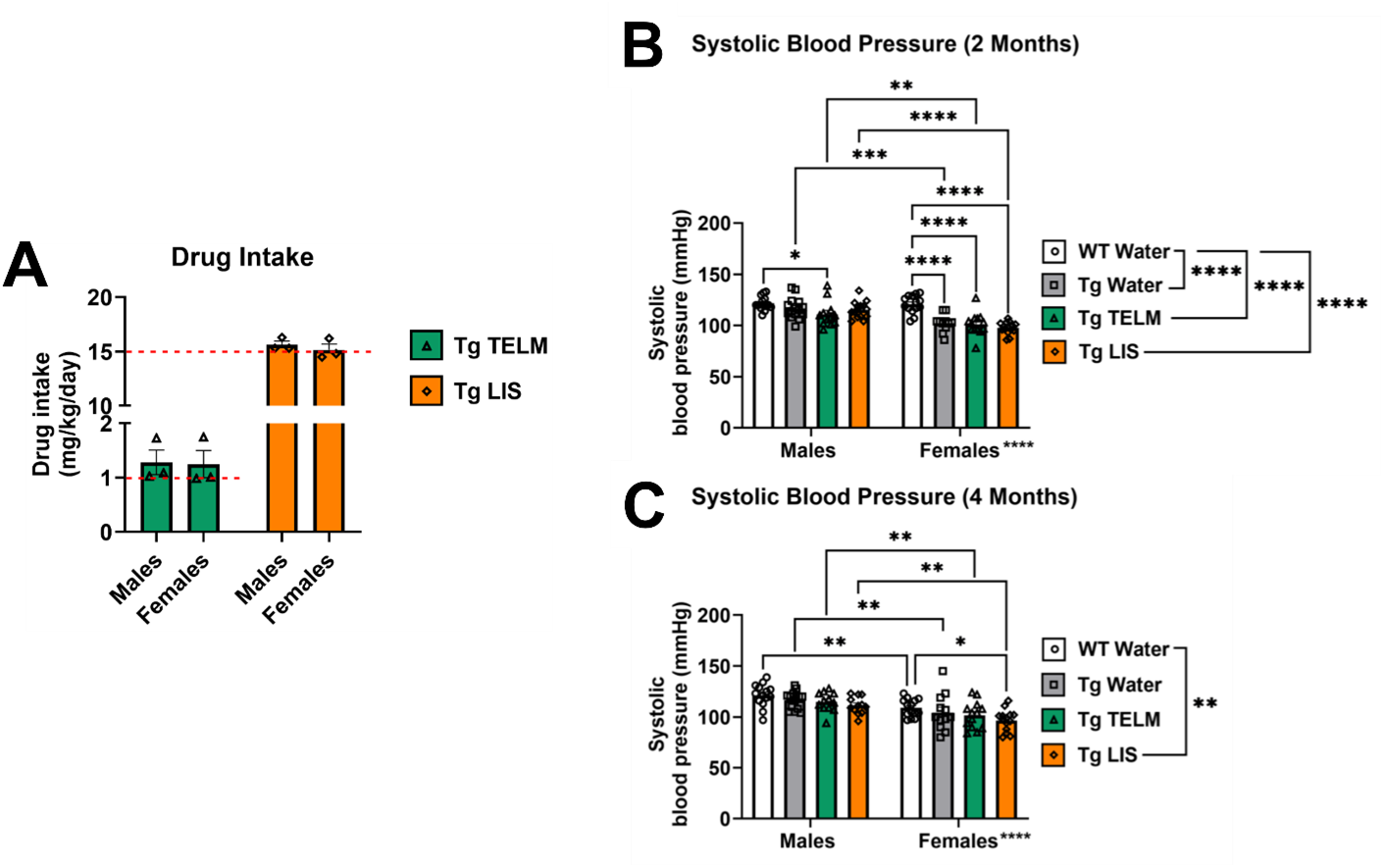
Drug intake and systolic BP. **(A)** There was no statistically significant difference observed between actual drug intake and the target doses of each drug (red dotted line). In the males, p=0.3333 for telmisartan and p=0.1978 for lisinopril. In the females, p=0.4228 for telmisartan and p=0.7805 for lisinopril. **(B)** Systolic blood pressure was measured using the tail cuff method at 2-months post-initiation of treatment. There was a main effect of group [F (3, 99) = 16.50, p<0.0001, η_p_^2^=0.3333]; overall, untreated Tg-SwDI (p<0.0001, d=1.187), telmisartan-treated Tg-SwDI (p<0.0001, d=1.474), and lisinopril-treated Tg-SwDI (p<0.0001, d=1.521) all had significantly lower systolic BP compared to WT mice groups. In the males, systolic BP of WTs was significantly higher than that of telmisartan-treated Tg-SwDI mice (p=0.0214, d=1.076). In the females, systolic BP of WTs was significantly higher than that of untreated Tg-SwDI mice (p<0.0001, d=2.148), telmisartan-treated Tg-SwDI mice (p<0.0001, d=1.972), and lisinopril-treated Tg-SwDI mice (p<0.0001, d=3.106). A statistically significant main effect of sex was observed [F (1, 99) = 36.50, p<0.0001, η_p_^2^=0.2694], with multiple comparisons showing significant sex differences within the untreated Tg-SwDI mice (p=0.0001, d=1.512), telmisartan-treated Tg-SwDI mice (p=0.0079, d=0.8411), and lisinopril-treated Tg-SwDI mice (p<0.0001, d=2.413). **(C)** At 4 months post-initiation of treatment, statistically significant main effect of group was observed [F (3, 98) = 4.568, p=0.0049, η_p_^2^=0.1227]; overall, lisinopril-treated Tg-SwDI mice had significantly lower systolic BP compared to WT mice groups (p=0.0025, d=0.9354). No statistically significant differences were observed between the male groups. In the females, lisinopril-treated Tg-SwDI mice had significantly lower systolic BP than WTs (p=0.0245, d=1.269). A statistically significant main effect of sex was observed [F (1, 98) = 35.62, p<0.0001, η_p_^2^=0.2665], with significant multiple comparison differences observed within the WTs (p=0.0044, d=1.246), untreated Tg-SwDI mice (p=0.0063, d=0.9053), telmisartan-treated Tg-SwDI mice (p=0.0044, d=1.135), and lisinopril-treated Tg-SwDI mice (p=0.0013, d=1.522).

### 3.2. No significant effect of drug treatment was observed in specific binding of ^125^I-SI Ang II in the brain of Tg-SwDI mice

A total of 13 brain regions were analyzed for ^125^I-SI Ang II binding: periventricular nucleus (PeVN), paraventricular nucleus (PVN), subfornical organ (SFO), median preoptic nucleus (MnPO), dorsomedial hypothalamus (DMH), vascular organ of lamina terminalis (OVLT), nucleus accumbens (NAcc), lateral septum, medial amygdala (MeA), central/basolateral amygdala (CeA/BLA), dorsal hippocampus (dHpc), sensorimotor cortex (SMC), and caudate putamen (CPu). For organizational purposes, analyses were divided into peri-hypothalamic regions [PeVN, PVN, SFO, MnPO, DMH, and OVLT (**Figure 5**)], regions of the striatum [NAcc and CPu (**Figure 6**)], regions of the amygdala [MeA and CeA/BLA (**Figure 7**)], lateral septum (**Figure 8**), hippocampus [dHpc (**Figure 9**)], and cortex [SMC (**Figure 10**)].

**Figure 5.**
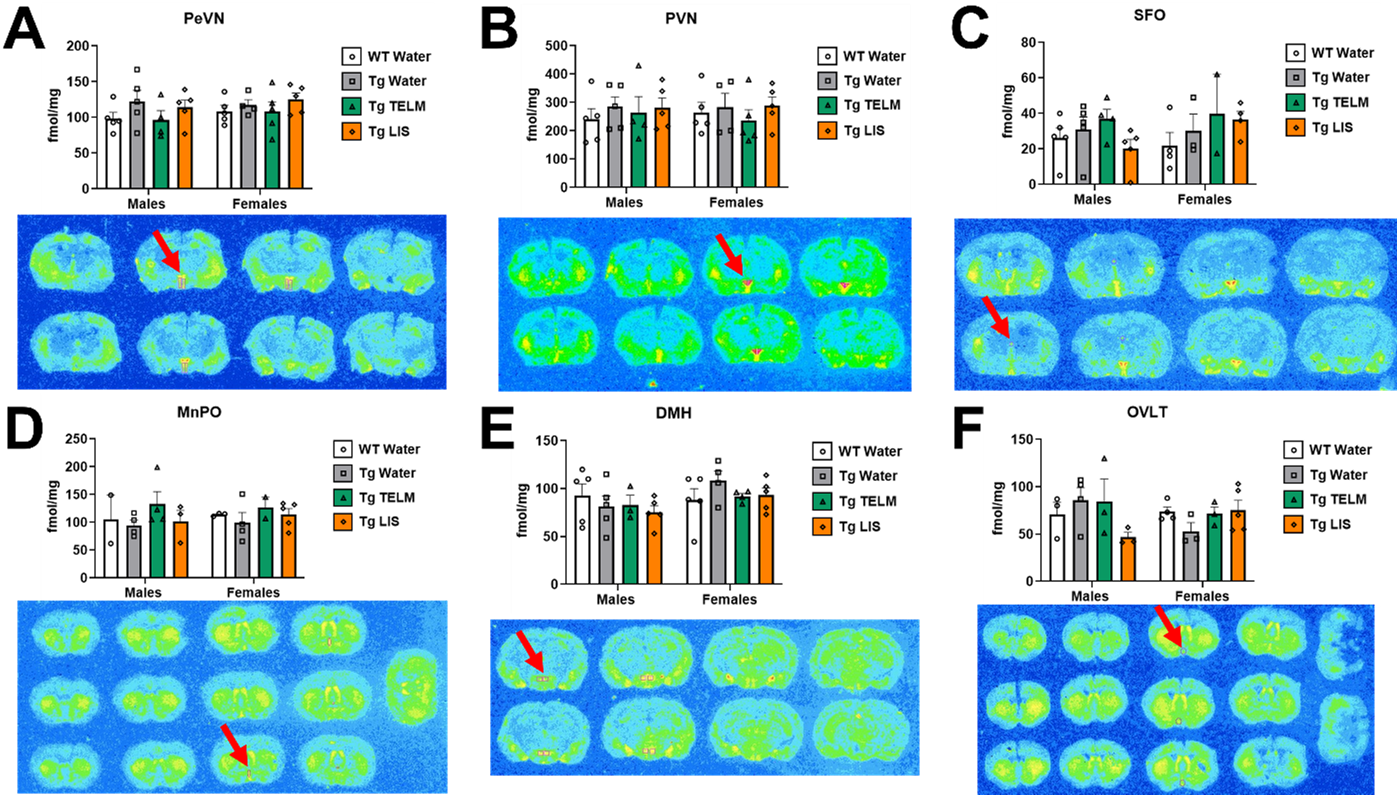
Specific binding of. ^125^**I-SI Ang II in the peri-hypothalamic regions of the brain of Tg-SwDI mice. (A)** A medium effect of treatment was observed in the PeVN, although not statistically significant [F (3, 30) = 1.538, p=0.2251, η_p_^2^=0.1333] (*N*=38; no outliers). Untreated Tg-SwDI mice showed elevated concentration of AT_1_R compared to WTs in both males (p=0.3966, d=0.8880) and females (p=0.9524, d=0.5083). In the males, telmisartan reduced specific binding in Tg-SwDI mice (p=0.3754, d=0.8593). No main effect of sex was observed [F (1, 30) = 0.799, p=0.379, η_p_^2^=0.026]. **(B)** No main effect of treatment was observed in the PVN [F (3, 30) = 0.4686, p=0.7064, η_p_^2^=0.0448] (*N*=38; no outliers). Untreated Tg-SwDI males showed elevated concentration of AT_1_R compared to WTs (p=0.8405, d=0.5506). In the females, telmisartan reduced specific binding in drug-treated Tg-SwDI mice (p=0.8463, d=0.5086). No main effect of sex was observed [F (1, 30) = 1.40E-04, p=0.991, η_p_^2^=4.67E-06]. **(C)** A medium effect of treatment was observed in the SFO, although not statistically significant [F (3, 24) = 1.123, p=0.3595, η_p_^2^=0.1231] (*N*=32; 4 outliers removed). Untreated Tg-SwDI females showed elevated concentration of AT_1_R compared to WTs (p=0.875, d=0.533). In the males, lisinopril reduced specific binding in drug-treated Tg-SwDI mice (p=0.6505, d=0.7766). In the females, lisinopril increased specific binding in drug-treated Tg-SwDI mice (p=0.9389, d=0.5023). No main effect of sex was observed [F (1, 24) = 0.4309, p=0.5178, η_p_^2^=0.0176]. **(D)** No main effect of treatment was observed in the MnPO [F (3, 19) = 1.070, p=0.3852, η_p_^2^=0.1446] (*N*=27; no outliers). Untreated Tg-SwDI females showed lower concentration of AT_1_R compared to WTs (p=0.939, d=0.504). Telmisartan increased specific binding, compared to untreated Tg-SwDI mice, in males (p=0.3646, d=1.122) and females (p=0.7840, d=0.7719). No main effect of sex was observed [F (1, 19) = 0.1261, p=0.7264, η_p_^2^=0.0066]. **(E)** No main effect of treatment was observed in the DMH [F (3, 28) = 0.3915, p=0.7601, η_p_^2^=0.0403] (*N*=36; 1 outlier removed). Untreated Tg-SwDI females showed elevated concentration of AT_1_R compared to WTs (p=0.5016, d=0.8193). Both drugs reduced specific binding, compared to untreated Tg-SwDI females (p=0.7085, d=1.042 for telmisartan and p=0.7242, d=0.7965 for lisinopril). A medium effect of sex was observed, although not statistically significant [F (1, 28) = 2.937, p=0.0976, η_p_ =0.0949]. **(F)** A medium effect of treatment was observed in the OVLT, although not statistically significant [F (3, 20) = 0.6746, p=0.5777, η_p_^2^=0.0919] (*N*=28; no outliers). Untreated Tg-SwDI males showed elevated concentration of AT_1_Rs compared to WTs (p=0.8116, d=0.5932). Lisinopril reduced specific binding compared to untreated Tg-SwDI males (p=0.1320, d=1.791). Untreated Tg-SwDI females showed reduced concentration of AT_1_Rs compared to WTs (p=0.6306, d=1.624). Both drugs increased specific binding, compared to untreated Tg-SwDI females (p=0.7464, d=1.296 for telmisartan and p=0.5198, d=1.090 for lisinopril). No main effect of sex was observed [F (1, 20) = 0.1747, p=0.6804, η_p_^2^=0.0087].

**Figure 6.**
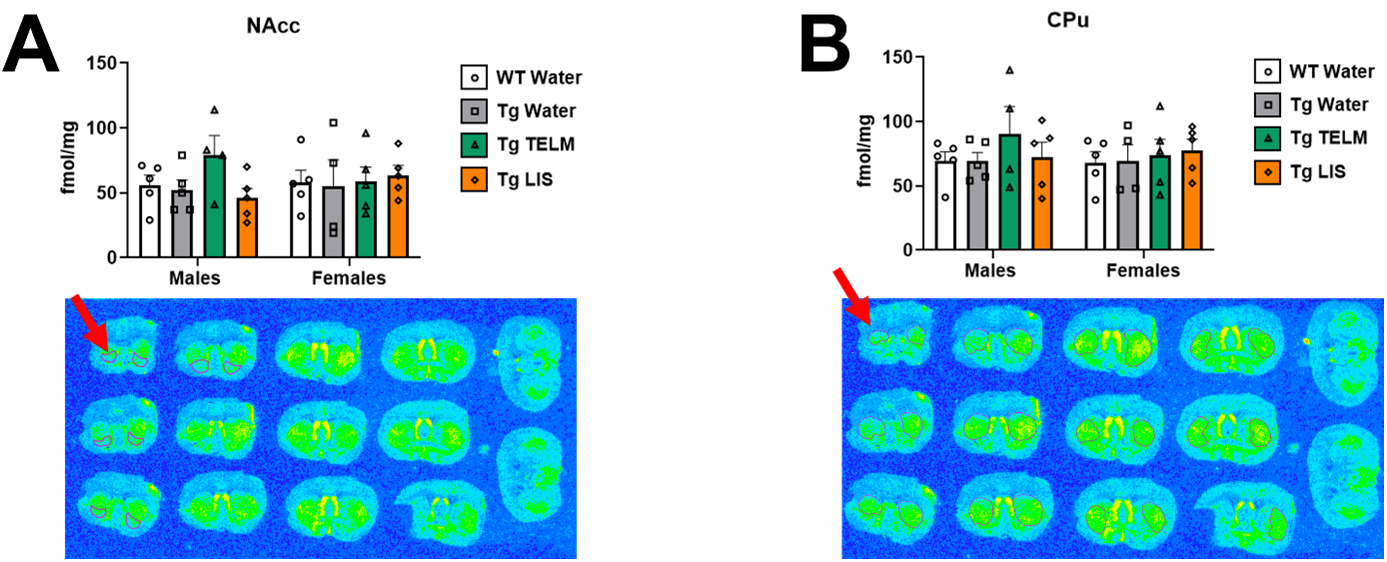
Specific binding of. ^125^**I-SI Ang II in the striatum of Tg-SwDI mice. (A)** A medium effect of treatment was observed in the NAcc, although not statistically significant [F (3, 30) = 0.8254, p=0.4903, η_p_^2^=0.0762] (*N*=38; no outliers). In the males, telmisartan increased specific binding compared to untreated Tg-SwDI mice (p=0.328, d=1.16). No main effect of sex was observed [F (1, 30) = 0.0040, p=0.9497, η_p_^2^=0.0001). **(B)** No main effect of treatment was observed in the CPu [F (3, 30) = 0.6130, p=0.6119, η_p_^2^=0.0578] (*N*=38; no outliers). In the males, telmisartan increased specific binding, compared to untreated Tg-SwDI males (p=0.5748, d=0.7090). No main effect of sex was observed [F (1, 30) = 0.149, p=0.702, η_p_^2^=0.005].

**Figure 7.**
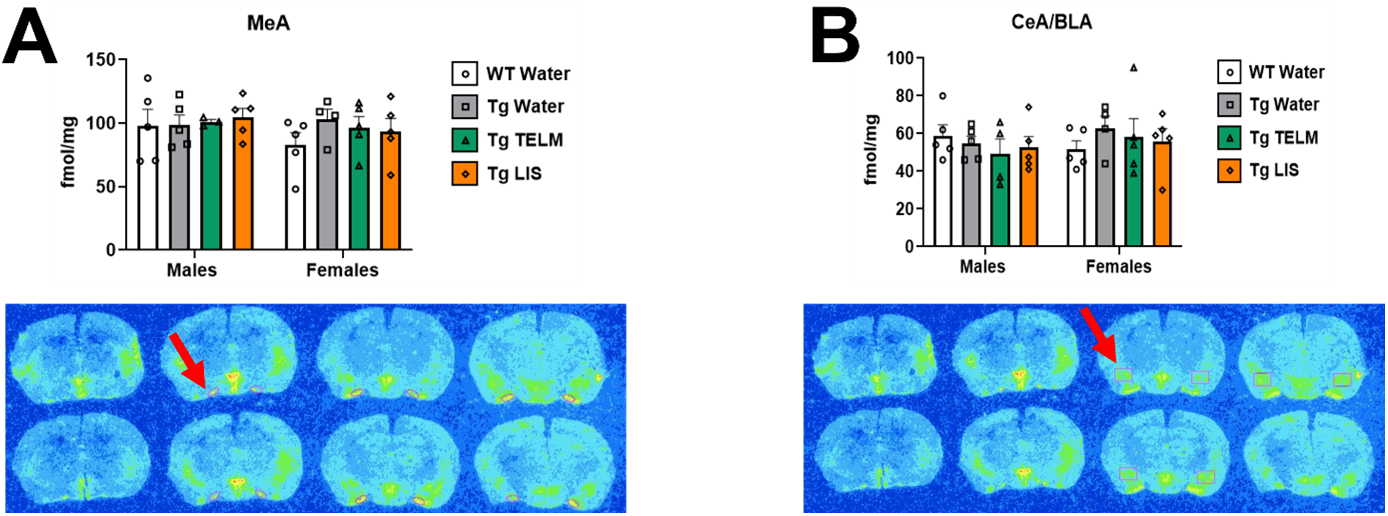
Specific binding of. ^125^**I-SI Ang II in the amygdala of Tg-SwDI mice. (A)** No main effect of treatment was observed in the MeA [F (3, 29) = 0.4964, p=0.6877, η_p_^2^=0.0488] (*N*=37; 1 outlier removed). Untreated Tg-SwDI females expressed elevated concentration of AT_1_Rs, compared to WTs (p=0.4747, d=1.018). No main effect of sex was observed [F (1, 29) = 0.9795, p=0.3305, η_p_^2^=0.0327]. **(B)** No main effect of treatment was observed in the CeA/BLA [F (3, 30) = 0.2351, p=0.8712, η_p_^2^=0.0230] (*N*=38; no outliers). Untreated Tg-SwDI females expressed elevated concentration of AT_1_Rs, compared to WTs (p=0.6727, d=0.9489). No main effect of sex was observed [F (1, 30) = 0.4594, p=0.5031, η_p_^2^=0.0151].

**Figure 8.**
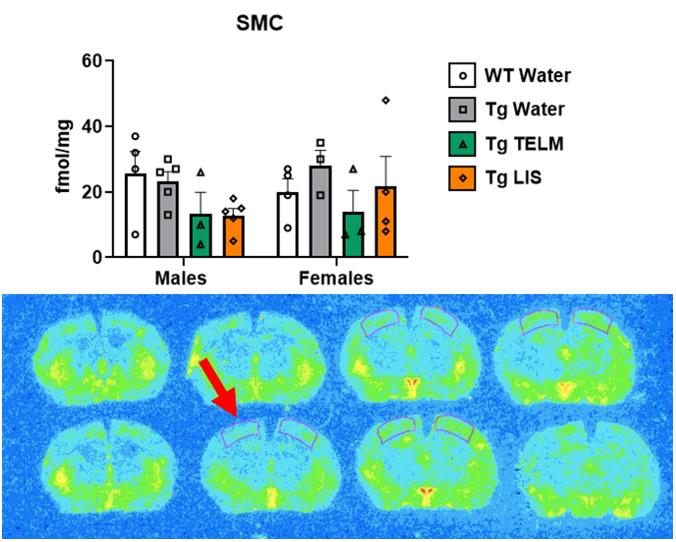
Specific binding of. ^125^**I-SI Ang II in the lateral septum of Tg-SwDI mice.** A medium effect of treatment was observed in the lateral septum, although not statistically significant [F (3, 29) = 0.7532, p=0.5295, η_p_^2^=0.0723] (*N*=37; 1 outlier removed). In the males, telmisartan increased specific binding, compared to untreated Tg-SwDI males (p=0.2965, d=1.777). Untreated Tg-SwDI females showed lower concentration of AT_1_Rs than WTs (p=0.7225, d=0.6548). No main effect of sex was observed [F (1, 29) = 0.0834, p=0.7748, η_p_^2^=0.0029].

**Figure 9.**
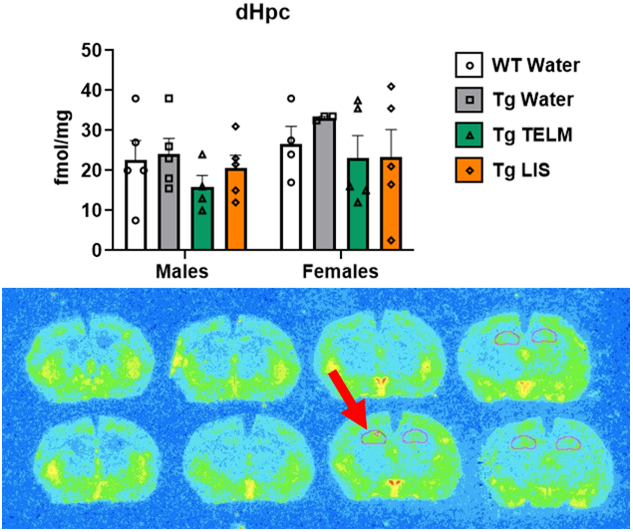
Specific binding of. ^125^**I-SI Ang II in the hippocampus of Tg-SwDI mice.** A medium effect of treatment was observed in the dHpc, although not statistically significant [F (3, 28) = 1.208, p=0.3252, η_p_^2^=0.1146] (*N*=36; 2 outliers removed). In the males, telmisartan decreased specific binding compared to untreated Tg-SwDI males (p=0.6194, d=1.081). Untreated Tg-SwDI females expressed slightly elevated concentration of AT_1_Rs compared to WTs (p=0.8351, d=0.9635). Both drugs decreased specific binding in females, compared to untreated Tg-SwDI females (p=0.5473, d=0.9962 for telmisartan and p=0.5555, d=0.7856 for lisinopril). A medium effect of sex was observed, although not statistically significant [F (1, 28) = 2.881, p=0.1007, η_p_^2^=0.0933].

**Figure 10.**
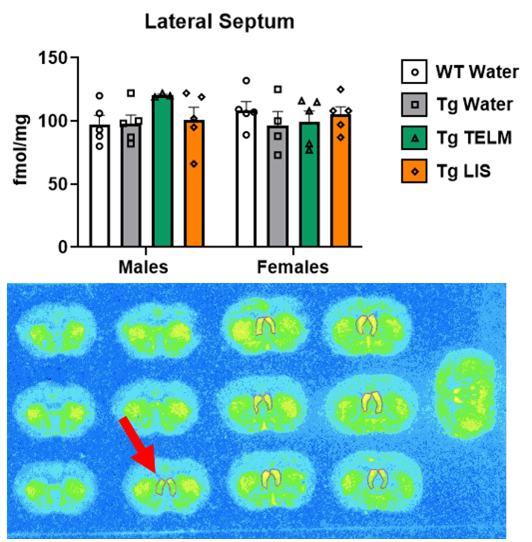
Specific binding of. ^125^**I-SI Ang II in the sensorimotor cortex of Tg-SwDI mice.** A large effect of treatment was observed in the SMC, although not statistically significant [F (3, 23) = 1.768, p=0.1814, η_p_^2^=0.1873] (*N*=31; 6 outliers removed). In the males, both drugs decreased specific binding, compared to untreated Tg-SwDI males (p=0.5952, d=1.151 for telmisartan and p=0.4333, d=1.766 for lisinopril). Untreated Tg-SwDI females expressed slightly elevated concentration of AT_1_Rs, compared to WTs (p=0.7629, d=0.9847). Telmisartan decreased specific binding in the females compared to untreated Tg-SwDI females (p=0.3974, d=1.421). No main effect of sex was observed [F (1, 23) = 0.3049, p=0.5862, η_p_^2^=0.0131].

Notably, there were no statistically significant differences in the specific binding of ^125^I-SI Ang II to AT_1_Rs between any groups across the brain regions measured. This arose from unexpectedly large variances, loss of some tissue sections in processing, and/or small sample sizes. However, statistical significance does not always equate to biological or clinical significance, as the relevance of an effect depends on its implications for physiology, disease progression, and/or treatment outcomes. Therefore, group differences with at least medium effect sizes (d ≥ 0.5, η_p_² ≥ 0.06) are discussed below. Medium effect sizes were defined as 0.5 ≤ d ≤ 0.799 and 0.06 ≤ η_p_ ^2^ ≤ 0.1399; while large effect sizes were defined as d ≥ 0.8 and η_p_ ^2^ ≥ 0.14.

Peri-hypothalamic regions: In the PeVN (**Figure 5A**), a medium effect of treatment was observed [F (3, 30) = 1.538, p=0.2251, η_p_ ^2^=0.1333]; overall, untreated (p=0.4639, d=0.7409, medium effect) and lisinopril-treated Tg-SwDI mice (p=0.4551, d=0.8106, large effect) had higher AT_1_R binding than WTs, while telmisartan reduced specific binding compared to untreated Tg-SwDI mice (p=0.4247, d=0.6608, medium effect). Untreated male (p=0.3966, d=0.8880, large effect) and female (p=0.9524, d=0.5083, medium effect) Tg-SwDI mice showed higher AT_1_R binding than WTs. Similarly, lisinopril-treated male (p=0.7420, d=0.7256, medium effect) and female (p=0.6958, d=0.8835, large effect) Tg-SwDI mice also showed higher AT_1_R binding than WTs. Additionally, telmisartan tended to decrease AT_1_R binding in Tg-SwDI males when compared to untreated Tg-SwDI mice (p=0.3754, d=0.8593, large effect). In the PVN (**Figure 5B**), untreated male Tg-SwDI mice showed higher AT_1_R binding than WTs (p=0.8405, d=0.5506, medium effect). The same was observed with lisinopril-treated Tg-SwDI males in comparison to WTs (p=0.8739, d=0.5004, medium effect). Additionally, telmisartan tended to decrease specific binding to the AT_1_R in Tg-SwDI females compared to untreated Tg-SwDI mice (p=0.8463, d=0.5086, medium effect). In the SFO (**Figure 5C**), a medium effect of treatment was observed [F (3, 24) = p=0.3595, η_p_^2^=0.1231]; overall, telmisartan-treated Tg-SwDI mice had higher AT_1_R binding than WTs (p=0.2934, d=0.9847, large effect) and lisinopril-treated Tg-SwDI mice (p=0.6008, d=0.6010, medium effect). Untreated female Tg-SwDI mice showed higher AT_1_R binding than WTs (p=0.8752, d=0.5326). The same was seen with telmisartan-(p=0.4947, d=0.8755, large effect) and lisinopril-treated (p=0.4930, d=1.171, large effect) Tg-SwDI females compared to WTs. Additionally, lisinopril reduced specific binding to the AT_1_Rs in the males (p=0.6505, d=0.7766, medium effect) and increased it in the females (p=0.9389, d=0.5023, medium effect) when compared to untreated Tg-SwDI mice. In the MnPO (**Figure 5D**), a large effect of treatment was observed [F (3, 19) = 1.070, p=0.3852, η_p_ ^2^=0.1446]; overall, telmisartan-treated Tg-SwDI mice had higher AT_1_R binding than WTs (p=0.7642, d=0.5811, medium effect) and untreated Tg-SwDI mice (p=0.3102, d=1.033, large effect), but lower than lisinopril-treated Tg-SwDI mice (p=0.6497, d=0.5895, medium effect). Untreated female Tg-SwDI mice showed lower AT_1_R binding than WTs (p=0.9389, d=0.503, medium effect). Meanwhile, telmisartan-treated male (p=0.7599, d=0.5630, medium effect) and female (p=0.9757, d=0.7753, medium effect) Tg-SwDI mice presented higher AT_1_R binding than WTs. Additionally, telmisartan increased specific binding to the AT_1_R in both male (p=0.3646, d=1.122, large effect) and female (p=0.7840, d=0.7719, medium effect) Tg-SwDI mice compared to untreated Tg-SwDI mice. In the DMH (**Figure 5E**), lisinopril-treated Tg-SwDI males showed lower AT_1_R binding than WTs (p0.5908, d=0.7527, medium effect); while untreated female Tg-SwDI mice showed higher AT_1_R binding than WTs (p=0.5016, d=0.8193, large effect). Additionally, telmisartan (p=0.7085. d=1.042, large effect) and lisinopril (p=0.7242, d=0.7965, medium effect) tended to reduce AT_1_R binding in Tg-SwDI females compared to untreated Tg-SwDI mice. A medium effect of sex was observed [F (1, 28) = 2.937, p=0.0976, η_p_^2^=0.0949]; overall, females had higher AT_1_R binding than males in the following groups: untreated Tg-SwDI mice (p=0.0685, d=1.156, large effect), telmisartan-treated Tg-SwDI mice (p=0.5877, d=0.7510, medium effect), and lisinopril-treated Tg-SwDI mice (p=0.1952, d=1.119, large effect). In the OVLT (**Figure 5F**), a medium effect of treatment was observed [F (3, 20) = 0.6746, p=0.5777, η_p_^2^=0.0919]; overall, lisinopril-treated Tg-SwDI mice showed lower AT_1_R binding than WTs (p=0.7889, d=0.5499, medium effect) and telmisartan-treated Tg-SwDI mice (p=0.5238, d=0.6037, medium effect). Untreated Tg-SwDI males showed higher AT_1_R binding than WTs (p=0.8116, d=0.5932, medium effect), while Tg-SwDI females showed lower (p=0.6306, d=1.624, large effect). In the males, lisinopril tended to reduce specific binding compared to WTs (p=0.5614, d=1.401, large effect) and to untreated Tg-SwDI males (p=0.1320, d=1.791, large effect). In the females, telmisartan (p=0.7464, d=1.296, large effect) and lisinopril (p=0.5198, d=1.090, large effect) tended to increase specific binding compared to untreated Tg-SwDI females.

Forebrain regions: In the NAcc (**Figure 6A**), a medium effect of treatment was observed [F (3, 30) = 0.8254, p=0.4903, η_p_^2^=0.0762]; overall, telmisartan-treated Tg-SwDI mice had higher AT_1_R binding than WTs (p=0.6811, d=0.5242, medium effect), untreated Tg-SwDI mice (p=0.5140, d=0.5608, medium effect), and lisinopril-treated Tg-SwDI mice (p=0.5625, d=0.5171, medium effect). When data was separated by sex, telmisartan-treated Tg-SwDI males showed higher AT_1_R binding than WTs (p=0.4652, d=0.9890, large effect), while lisinopril-treated Tg-SwDI mice showed lower (p=0.9067, d=0.5880, medium effect). Additionally, telmisartan also increased specific binding to the AT_1_R in males compared to untreated Tg-SwDI males (p=0.3276, d=1.156, large effect). In the CPu (**Figure 6B**), telmisartan-treated Tg-SwDI males (p=0.5673, d=0.7041, medium effect) and lisinopril-treated females (p=0.9237, d=0.5057, medium effect) showed higher AT_1_R binding than WTs of the same sex. Additionally, telmisartan increased specific binding to the AT_1_R in males compared to untreated Tg-SwDI males (p=0.5748, d=0.7090, medium effect).

Amygdala regions: In the MeA (**Figure 7A**), untreated (p=0.4747, d=1.018, large effect) and telmisartan-treated (p=0.7161, d=0.6604, medium effect) Tg-SwDI females showed higher AT_1_R binding than WTs. In the CeA/BLA (**Figure 7B**), telmisartan-treated Tg-SwDI males showed lower AT_1_R binding than WTs (p=0.7468, d=0.6688, medium effect); while untreated Tg-SwDI females showed higher (p=0.6727, d=0.9489, large effect).

Lateral septum (**Figure 8**), dHpc (**Figure 9**), and SMC (**Figure 10**): In the lateral septum, a medium effect of treatment was observed [F (3, 29) = 0.7532, p=0.5295, η_p_^2^=0.0723]; overall, telmisartan-treated Tg-SwDI mice had higher AT_1_R binding than untreated Tg-SwDI mice (p=0.4487, d=0.7324, medium effect). In the males, telmisartan-treated Tg-SwDI mice showed higher AT_1_R binding than WTs (p=0.2820, d=1.774, large effect) and untreated Tg-SwDI males (p=0.2965, d=1.777, large effect). In the females, untreated (p=0.7225, d=0.6548, medium effect) and telmisartan-treated (p=0.8410, d=0.5264, medium effect) Tg-SwDI mice showed lower AT_1_R binding than WTs. In the dHpc, a medium effect of treatment was observed [F (3, 28) = 1.208, p=0.3252, η_p_^2^=0.1146]; overall, telmisartan-treated mice had lower AT_1_R binding than WTs (p=0.7210, d=0.5083, medium effect) and untreated Tg-SwDI mice (p=0.2895, d=0.9842, large effect), while lisinopril-treated mice had lower AT_1_R binding than untreated Tg-SwDI mice (p=0.5299, d=0.6627, medium effect). In the males, telmisartan-treated Tg-SwDI mice showed lower AT_1_R binding than WTs (p=0.7582, d=0.7250, medium effect). Untreated Tg-SwDI females showed higher AT_1_R binding than WTs (p=0.8351, d=0.9635, large effect). Additionally, telmisartan reduced specific binding in both males (p=0.6194, d=1.081, large effect) and females (p=0.5473, d=0.9962, large effect) compared to untreated Tg-SwDI mice; while lisinopril reduced specific binding in females only (p=0.5555, d=0.786, medium effect). A medium effect of sex was observed [F (1, 28) = 2.881, p=0.1007, η_p_^2^=0.0933]; overall, female mice had higher AT_1_R binding than males of the following groups: untreated Tg-SwDI mice (p=0.2337, d=1.262, large effect) and telmisartan-treated Tg-SwDI mice (p=0.2855, d=0.7402, medium effect). In the SMC, a large effect of treatment was observed [F (3, 23) = 1.768, p=0.1814, η_p_^2^=0.1873]; overall, telmisartan-treated Tg-SwDI mice had lower AT_1_R binding than WTs (p=0.4022, d=0.8868, large effect), while untreated Tg-SwDI mice had higher AT_1_R binding than telmisartan-(p=0.2045, d=1.398, large effect) and lisinopril-treated (p=0.4154, d=0.9375, large effect) Tg-SwDI mice. In the males, telmisartan-(p=0.4431, d=0.9961, large effect) and lisinopril-treated (p=0.2974, d=1.383, large effect) Tg-SwDI mice showed lower AT_1_R binding than WTs. In the females, untreated Tg-SwDI mice showed higher AT_1_R binding than WTs (p=0.7629, d=0.9847, large effect); while telmisartan-treated Tg-SwDI mice showed lower (p=0.8825, d=0.6325, medium effect). Additionally, telmisartan reduced specific binding in both males (p=0.5952, d=1.151, large effect) and females (p=0.3974, d=1.421, large effect) compared to untreated Tg-SwDI mice; while lisinopril reduced specific binding in males only (p=0.4333, d=1.766, large effect).

## 4. Discussion

Previous studies have shown that AD is linked to increased expression of components of the RAAS, including ACE, Ang II, and AT_1_Rs, especially in the hippocampus, frontal cortex, and caudate nucleus^19, 68–70^, leading to overactivation of the classical RAAS arm. Similarly, in animal models of AD, increased expression of astrocytic AT_1_R protein was detected in the hippocampus^71^. However, this is the first study to assess AT_1_Rs in postmortem CAA brains, across multiple regions, and to determine if RAAS-targeting drugs can affect AT_1_R expression in the brains of CAA-prone mice. Tg-SwDI mice that exhibit CAA were treated with a sub-depressor dose of either telmisartan or lisinopril for 5 months. Postmortem, RAR was performed to determine levels of AT_1_R capable of binding an Ang II ligand in 13 brain regions, with varying levels of AT_1_R expression. The results were compared to untreated Tg-SwDI mice, as well as WT mice. While there have been studies of AT_1_R-like protein using immunohistochemical methods in AD mice^19^, this methodology lacks specificity^72–77^. Additionally, the antibodies do not distinguish functional from non-functional protein or protein fragments. The lack of AT_1_R antibody specificity is equally applicable to Western blots. Results showed no statistically significant differences among groups with or without drug treatment in any of the 13 regions analyzed. Trends with medium to large effect sizes were observed; however, there were no systematic increases or decreases in AT_1_R expression across the brain regions analyzed.

Circulating Ang II does not usually penetrate the BBB in a healthy brain, due to its size and hydrophilicity, although local production in the brain does occur^78, 79^. Additionally, several circumventricular organs, e.g., SFO, OVLT, and area postrema of the brain, are enriched in AT1Rs that can respond to bloodborne Ang II^80–84^. In hypertension and inflammatory brain diseases, such as stroke, dementia, and TBI, where disruption of the BBB occurs, more Ang II is capable of penetrating through the barrier, causing overactivation of the classical RAAS arm^78, 85–87^. Similarly, it has also been shown that ACE, the rate-limiting enzyme in the formation of Ang II is also elevated in conditions such as AD^88^. The overactivation of AT_1_Rs by Ang II results in neurotoxicity, neuroinflammation, oxidative stress, and Aβ deposits, as well as worse cognitive performance and onset of dementia^19, 85^. Therefore, centrally-acting RAAS-targeting drugs that block AT_1_Rs (ARBs) or prevent the formation of Ang II (ACE inhibitor) in these conditions can have neuroprotective effects^86^.

The first group of RAR analyses included peri-hypothalamic regions, with high levels of AT_1_R expression^84^. The hypothalamus includes several distinct nuclei responsible for maintaining homeostasis by integrating sensory inputs and activating autonomic, endocrine, and behavioral responses^89, 90^. Under normal conditions, Ang II is not able to cross the BBB to act on AT_1_Rs present in certain regions of the hypothalamus^78, 85^, although the SFO, which is a sensory circumventricular organ, as noted above, is able to sense circulating levels of Ang II to transmit the information to autonomic centers in the hypothalamus, such as the PVN^78^, which is considered a major center for mediating central ang II activity, by stimulating sympathetic activities^91^. It has been shown that blocking AT_1_Rs in these regions can reduce sympathetic activity^91^.

Subdivisions of the peri-hypothalamic regions that were analyzed in this study include the PeVN (involved in hormone secretion^92, 93^), PVN (involved in hormone secretion, homeostasis, cardiovascular and immune functions, and reproduction^94, 95^), MnPO (involved in body fluid balance, temperature regulation, and cardiovascular and immune functions^96, 97^), SFO (circumventricular organ involved in homeostasis and cardiovascular and immune response regulation^98, 99^), OVLT (circumventricular organ involved in osmoregulation^100^), and DMH (involved in sleep, body temperature, and cardiovascular response to stress^101–103^). In general, untreated Tg-SwDI mice tended to have higher AT_1_R binding compared to WT mice with some sex-specific findings, including PeVN, PVN (males), DMH (females), SFO (females), and OVLT (males). A different trend was seen in the MnPO and OVLT, with Tg-SwDI mice having lower AT_1_R binding, though these findings were specific to females. Treatment with RAAS-targeting drugs tended to normalize AT_1_R binding levels in Tg-SwDI mice, with effects depending on drug, region, and sex. Telmisartan reduced AT1R in PeVN, PVN (males), DMH (females), and OVLT (females), suggesting a protective effect; however, it increased AT_1_R in MnPO and SFO, even in regions where AT_1_R binding was already augmented in untreated Tg-SwDI mice. Lisinopril had some benefits, but less widespread effects, normalizing AT_1_R in the DMH (females) and OVLT (both sexes); however, there was overshoot in the OVLT of males.

The second group of RAR analyses included regions of the striatum, an area involved in decision making functions (e.g., motor control and emotions) ^104^, with moderately high levels of AT_1_R expression^105^. In the striatum, AT_1_Rs regulate dopamine signaling^106^. Dopamine contributes to sensorimotor coordination and reward-related stimuli in the striatum^107^. In this study, in both the NAcc (the brain’s reward center^108^) and the CPu (involved in motor control, cognition, and emotions^109, 110^), AT_1_R levels were similar between Tg-SwDI and WT mice, with no sex-specific alterations and no effect of drug treatment.

The third group of RAR analyses included regions of the amygdala, a brain region that is responsive to AT_1_R stimulation^111^. Part of the limbic system, the amygdala is responsible for processing emotions, controlling behavior, and memory formation^112^. Activation of AT_1_Rs in the amygdala has been associated with fear and threat-learning and memory, as well as anxiety-like behaviors^113^. In the MeA (involved in processing social and non-social behaviors^114^) and CeA/BLA (involved in emotion processing^115^), Tg-SwDI females had higher AT_1_R levels than WTs, while no differences were seen between male Tg-SwDI and WTs. Additionally, while not significant, drug-treated Tg-SwDI females tended to have AT_1_R binding levels closer to that of WT controls.

In the lateral septum (region that mediates aggressive behavior^116^), which expresses moderately high levels of AT_1_Rs and AT_2_Rs^117^, sex-specific differences in AT_1_R expression were seen in Tg-SwDI mice; Tg-SwDI females expressed lower AT_1_R levels than WTs, while AT_1_R expression in Tg-SwDI males did not differ from WTs. In the males only, telmisartan increased AT_1_R levels compared to untreated Tg-SwDI male mice.

In the dHpc (involved in spatial memory, learning, and navigation^118, 119^) and SMC (involved in integrating sensory information with motor function^120^), AT_1_R expression occurred at low levels. Nevertheless, AT_1_R expression was increased in Tg-SwDI female mice only compared to female WTs. In the dHpc, AT_1_R levels were normalized in Tg-SwDI females treated with either lisinopril or telmisartan, while in the SMC, AT_1_R levels were normalized by telmisartan treatment, only. In males, drug treatment tended to reduce AT_1_R binding, even though they were not altered in Tg-SwDI compared to WT mice.

As mentioned previously, it has been established that AD brains present elevated levels of ACE, Ang II, and AT_1_Rs^19, 68–70^. While previous studies have investigated AT_1_R levels in the cortex of humans^19, 68^ and hippocampus of AD mouse models^71^, they used antibody-based techniques (i.e. immunohistochemistry, Western blot), which, as noted above, lack specificity. Although BBB disruption does occur in CAA, that is only observed in late disease stage^121–123^. It is possible that the age of our mice at euthanasia was not late enough into the disease to observe significant impairments to the BBB, which would explain lack of significant differences between Tg-SwDI mice and WTs. Additionally, in AD, pathology is first observed in the entorhinal cortex, followed by the hippocampus, thalamus, and eventually the neocortex^19, 78, 87, 124^. As AD pathology spreads to these brain regions, worsening cognition is observed, progressing from mild cognitive impairment to severe dementia^124^. In contrast, CAA pathology in the Tg-SwDI moue model used here is mainly observed in the hippocampus, followed by the thalamus, and is almost absent in the cortex^54^. However, when it comes to AT_1_R dysregulation, we saw no statistically significant differences in brain regions between Tg-SwDI mice and WTs, although trends with medium to large effect sizes were observed. In most but not in all brain regions analyzed, Tg-SwDI females expressed higher levels of AT_1_R than their WT counterparts, while the same was not observed in males to the same magnitude. Although this has not been previously shown, it is on par with sex-difference findings that show that Tg-SwDI females have higher number of cerebral microbleeds, more severe cognitive impairment, and lower levels of neural repair cytokines than Tg-SwDI males^125^, meaning therefore, that Tg-SwDI females have more severe CAA-related neuropathology and symptoms than Tg-SwDI males.

It has been reported that, although RAS-targeting drugs reduce Ang II formation (ACE inhibitors) or prevent its effects by blocking the AT_1_R (ARBs), a compensatory upregulation of AT_1_Rs can occur to maintain a balance of the signaling cascade^126^. This could explain why AT_1_R binding/expression was increased by drug treatment in some brain regions. However, further exploration is needed to determine if these changes are physiologically significant. On the other hand, there are reports of AT_1_R down-regulation in response to chronic losartan treatment in the hamster heart, although the ACE inhibitor quinapril was without such effect^127^, which could potentially explain the lower expression of AT_1_Rs in some brain regions of the telmisartan treated

Tg-SwDI mice. Additionally, despite substantial effect sizes being observed, none of them attained statistical significance. This could be due to a variety of factors, such as unexpected large variances, loss of some tissue sections in processing, and small sample sizes due to attrition over the 5-month course of the study.

## 5. Conclusions

Although we hypothesized that drug treatment would prevent differences in brain AT_1_R expression in Tg-SwDI mice associated with CAA, results from RAR did not support this, as AT_1_R levels in transgenic animals were not significantly different from that of WTs, and drug treatment did not significantly alter AT_1_R levels in the Tg-SwDI mice. We did, however, observe a medium effect of sex in the DMH and dHpc regions, with females expressing higher AT_1_R expression. Further studies with increased sample sizes would be required to determine if the trends observed are pathophysiological and pharmacologically significant.

## Supporting information

Figure 1 - Publication License

Figure 2 - Publication License

Figure 3 - Publication License

## Acknowledgments

The authors would like to thank Yazmin Restrepo, Ariana Hernandez, Chana Vogel, Victoria Pulido-Correa, Kaitlin Martin, Shay Moen, and Julianna Bonetti for their help conducting experiments and analyses.

