## Supplementary material for "Chronic Treatment of a Mouse Model of Cerebral Amyloid Angiopathy and Brain AT_1_ Receptor Expression": Figure 3 - Publication License

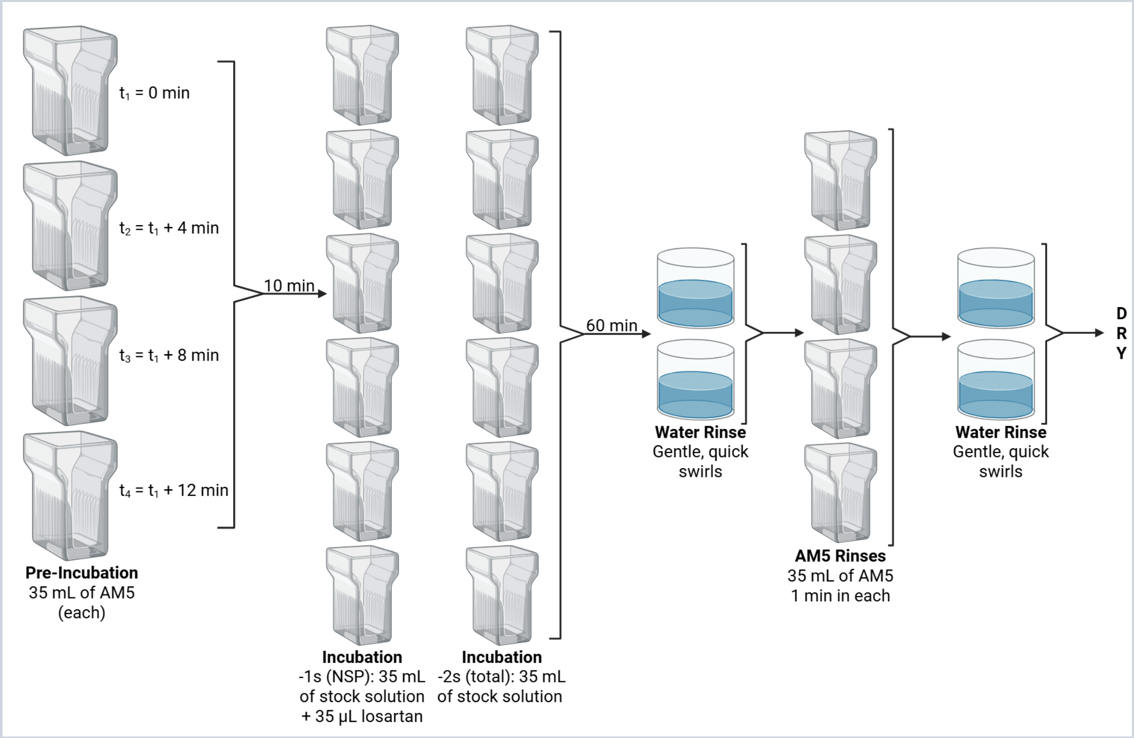


*For any questions regarding this document, or other questions about publishing with BioRender, please refer to our BioRender Publication Guide, or contact BioRender Support at.*
